# Flight feathers evolve fastest near the tip of the wing

**DOI:** 10.1101/2025.04.17.649363

**Authors:** Jonathan A. Rader, Eli B. Bradley, Mary E. Petersen, Daniel R. Matute

## Abstract

Form-function relationships are foundational to evolutionary biology, but we are only beginning to understand the mechanisms that link biomechanical function to morphological evolution. In bird wings, the pace of morphological evolution follows a gradient of aerodynamic force production along the wing, with the fastest rates experienced at the wingtip. We examined how this gradient has manifested in the evolution of the individual flight feathers that comprise the wing surface. Specifically, we studied whether the flight feathers show a pattern of allometric scaling with respect to avian body size, and if the allometric pattern varies along the wing. We also asked whether the pace of evolution of the primary flight feathers follows the same gradient pattern as the overall wing morphology, and if there is modularity within the feather series. We measured the lengths of each of the primary flight feathers from 509 wings representing 514 bird species and calculated morphological disparity and evolutionary rate for each feather in the primary sequence. We found that evolutionary rate and disparity increase significantly toward the wingtip. We also found evidence for modularity within the feather sequence, but we were unable to positively delineate those modules. Our study provides further evidence that the mechanical sensitivity of morphological traits predicts their evolution, with highly sensitive features biased toward higher rates, and that it scales across multiple levels of anatomical organization.

## Introduction

Form-function relationships are fundamental to morphological evolution because they dictate how structural adaptations enhance an organism’s ability to survive and reproduce within its environment (Bock and von Wahlert 1965; Lauder 1990; Irschick 2002; Carroll 2005). Inter and intraspecific differences in shape and size allow animals to access and compete for resources in different ways (Schluter 2000; Pfennig and Pfennig 2009). Natural selection favors trait variants that optimize functions such as locomotion, feeding, and defense. Ultimately, the map between morphology and function determines the adaptive potential of a genotype, guiding evolutionary trajectories, and enabling the exploitation of ecological niches (Higham et al. 2021). Even though form-function relationships are a cornerstone of evolutionary biology, we are only beginning to understand the evolutionary mechanisms that link biomechanical function to morphological evolution (Muñoz et al. 2018; Muñoz and Price 2019). Birds are diverse in their morphology, behavior, and ecology (Barnagaud et al. 2012; Jetz et al. 2012; Barrowclough et al. 2016; Felice et al. 2019), and have provided some of the clearest examples of the relationship between morphology and fitness. The rapid evolution of cranial morphology among the species of Darwin’s finches followed changes in diet type and resource availability (Schluter and Grant 1984). Similarly, prey type in raptors has a larger impact on talon evolution than body size (Tsang et al. 2019).

Wings are complex structures that serve a variety of functions among birds (del Hoyo et al. 1992; Sibley et al. 2009), especially the production of aerodynamic forces for flight. Flight forces are generated through the fluid-structure interaction of the wings with the air (or water) through which the birds are moving (Vogel 1981), and the magnitude of those forces and efficiency with which they are produced is related to their form (Vogel 1981; Shyy et al. 2007, 2010). Bird wings include skeletal, muscular, and integumentary elements, including the feathers that form the external morphology and aerodynamic surface of the wing (Figure 1). Wing shape corresponds with flight style, migratory habits, and other key aspects of avian biology (e.g., Lockwood et al. 1998; Claramunt et al. 2012; Stoddard et al. 2017; Sheard et al. 2020), and it is strongly predicted by phylogenetic relatedness (Baliga et al. 2019). The aerodynamic surface of the wing is composed of the primary, secondary and tertial flight feathers, along with the coverts, all of which vary in their size and shape along the length of the wing (Worcester 1996; Wang et al. 2012; Wang and Clarke 2015). Primary feathers are those that originate on the carpometacarpus (the bones of the avian hand), while the secondaries emerge from the ulna (Figure 1). The tertials bridge the gap between the secondary feathers and the body, and the coverts form the leading edge of the wing and cover the insertion of the flight feathers into the arm.

**Figure 1.**
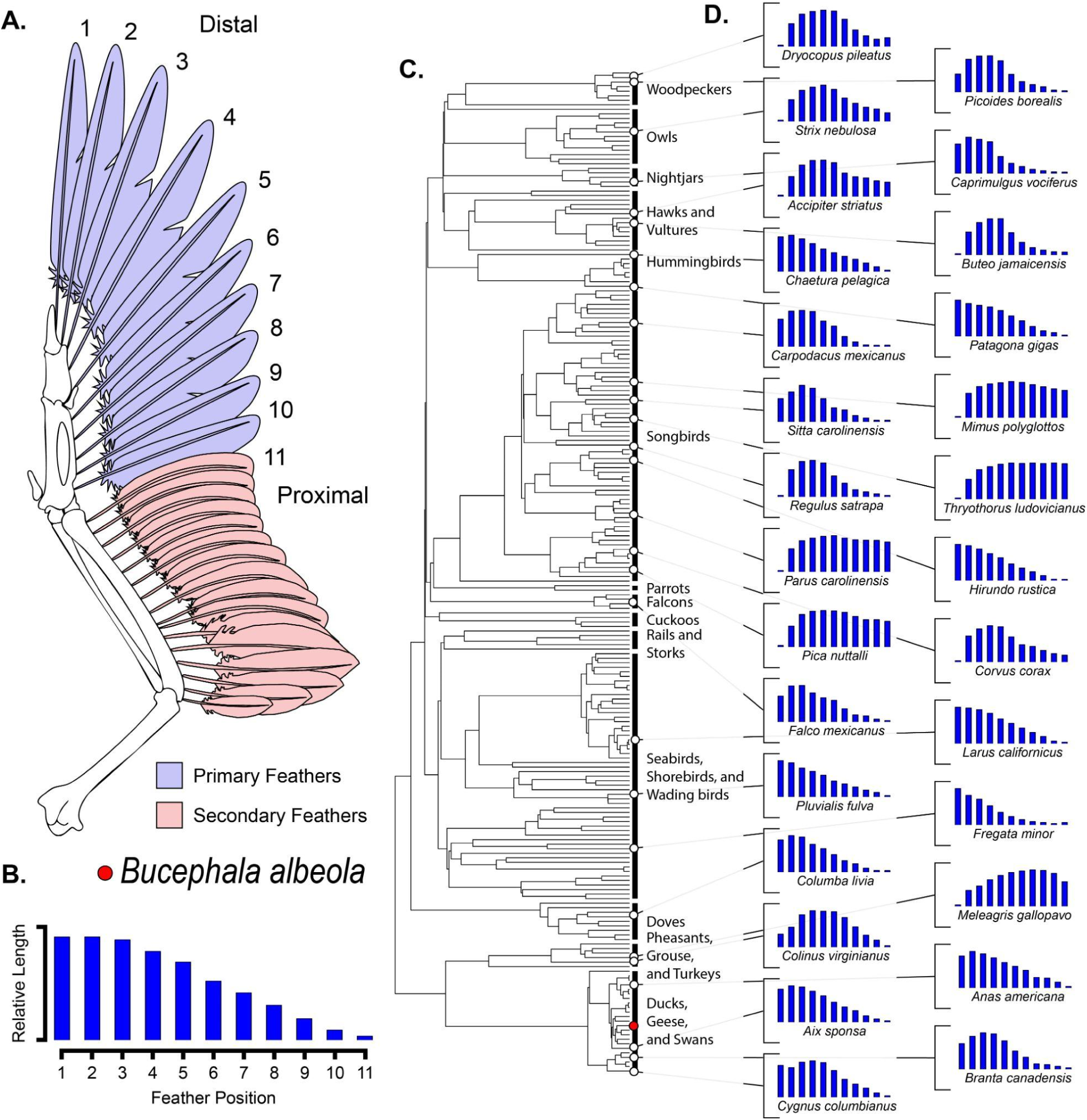
Wing diagram and phylogeny depicting taxa and wing portion included in this study. A) Diagram of the skeleton and the flight feathers. Primary feathers, shown in blue, are the flight feathers associated with the hand portion of the wing (handwing), and the secondary feathers, shown in pink, are associated with the ulna and form a portion of the armwing. We measured the eleven distal-most feathers from the wings (labeled 1 to 11 from distal to proximal), as their relative lengths capture the shape of the wingtip. The example wing shows the feather lengths from a species of duck, the Bufflehead (*Bucephala albeola*). B) The relative lengths of the 11 measured feathers is shown by the blue bars. C) The phylogeny shows the taxonomic breadth included in our sample of 214 species of birds (the full species list can be found in the supplemental data file). D) Subplots, similar to Panel B, show the relative lengths of feathers for 28 species. Open circles at the phylogeny tips show the position of each example species within the phylogeny, and are linked to each subpanel by lines. The red circle shows the position of *Bucephala albeola* from Panel B.

The primary flight feathers contribute to overall wing length, particularly in the handwing at the distal end of the series. Proximal primary feathers, meanwhile, merge with the secondary feathers, forming part of the wing chord along with the leading-edge anatomy (see Figure 1A). The number of primaries differs among bird taxa, ranging from 9 in in some passerines (though most have 10) to 12 in storks, flamingos, and grebes (del Hoyo et al. 1992; Sibley et al. 2009; Jenni and Winkler 2020). Combined, the primaries define the area of the handwing, and their relative lengths determine the shape of the wingtip (Kipp 1959; Lockwood et al. 1998). The length of the secondary flight feathers correlates with wing chord (Lockwood et al. 1998), while the number of secondaries varies among taxa and is associated with overall wing length, particularly in the armwing (Sibley et al. 2009). Both primary and secondary feathers play essential roles in flight, but primaries have received special attention due to their role in shaping the wingtip—an aerodynamically critical region (Shyy et al. 2010). Much attention has been paid to the relative lengths of the primary feathers (including wingtip shape indices, e.g., Kipp 1959; Lockwood et al. 1998), the diversity of wingtip geometry, and its correlates in other aspects of bird biology (e.g., Mulvihill and Chandler 1990; Lockwood et al. 1998; Stoddard et al. 2017; Sheard et al. 2019, 2020; Pigot et al. 2020, and many others). Meanwhile, the allometric scaling of the individual feathers with respect to body size and the tempo and mode of their evolution has remained understudied. To bridge this gap, we investigated whether the evolution of the flight feathers, particularly of the primaries, is driven by flight physics.

Flapping flight is characterized by rotation of the wing around a pivot point at the shoulder. The rotational motion generates an increasing proximo-distal velocity gradient along the length of the wing (Weis-Fogh 1972). Because physical dynamics such as aerodynamic forces (lift and drag) and inertial moment both vary as a square-function of velocity, the velocity gradient in the wing also generates force gradients along the wing (Weis-Fogh 1972; van den Berg and Rayner 1995). The model predicting these spanwise gradients is based on a simple two-dimensional representation of the wing, however they are also influenced by three-dimensional induced flow and rotational lift effects. The generation of tip vortices creates pressure differences along the wing that interact with leading-edge vortices, contributing to a net increase in the force per unit area along the length of the wing during flapping flight (Birch and Dickinson 2001; Shyy et al. 2007, 2010), although wingtip vortices actually reduce aerodynamic forces near the tip of the wing while gliding (Shyy et al. 2007). These effects are partially mitigated by the tapering of wing chord and mass from wing root-to-tip, a balance which is expected to differ among taxa with differing wing geometries. Overall, though, despite the influences of vorticity and wing taper, distal portions of the wing experience greater aerodynamic forces (Weis-Fogh 1972; Shyy et al. 2010) and inertial moments (van den Berg and Rayner 1995) than proximal ones, meaning that evolutionary changes in shape have increasingly pronounced effects on the physical dynamics of the wing the further they occur from the wing root.

Traits that have strong influence on the functional output of a biomechanical system may be under strong selective pressure, in turn leading to increased rates of phenotypic evolution (Anderson and Patek 2015; Muñoz et al. 2017, 2018). For example, in four-bar linkage systems such as those in the jaws of bony fish (Muller 1987; Westneat 1990), the links that have the greatest impact on kinematic transfer also have the highest evolutionary rates (Anderson and Patek 2015; Muñoz et al. 2017, 2018). In bird wings, the evolutionary rate of aerodynamically-relevant traits such as wing chord and camber increase in a gradient from the base of the wing toward its tip (Rader and Hedrick 2023; see also Eliason et al. 2023), following a similar pattern to inertial moment and the production of aerodynamic forces during flapping flight (Weis-Fogh 1972; van den Berg and Rayner 1995). The flight feathers are a set of serially homologous elements arranged spanwise along the wing that are superimposed upon the root-to-tip force gradient of aerodynamic forces (Weis-Fogh 1972). Thus, we predicted that because the relative lengths of the primary feathers determine wingtip geometry (see Figure 1), the base-to-tip gradient would also be apparent in the evolutionary dynamics (tempo and mode; Simpson 1944) of primary feather lengths. Specifically, we predicted that evolutionary rate and morphological disparity would increase progressively from proximal to distal across the sampled feather sequence, and that the proximal feathers would show relatively greater evolutionary conservatism than more distal ones.

The flight feathers are discrete anatomical units (Wessells 1965; Prum 1999; Busby et al. 2020) arranged in a series along the length of the wing, with continuous force and possibly selective gradients superimposed across them (Weis-Fogh 1972; van den Berg and Rayner 1995; Shyy et al. 2010). Each feather might therefore be exposed to their own selection regime and thus evolve independently from each other. On the other hand, the shape of each feather contributes to the overall geometry of the wing and its aerodynamic output (Lockwood et al. 1998; Swaddle and Lockwood 2003), and because of their shared development (Busby et al. 2020), the primary feathers might evolve as an integrated unit. We therefore tested two competing hypotheses regarding the morphological integration of the distal flight feathers: 1) The feathers are discrete anatomical and evolutionary units, and do not show any organization into modules. 2) The feathers show regional modularity along the length of the wing into groups of feathers that share similar evolutionary dynamics (allometry, evolutionary tempo and mode).

Allometric scaling refers to the changes in the relative size of morphological traits across a range of body size, and can refer to either changes in trait proportions through ontogeny or comparatively across taxa. In this study, we are focused on evolutionary allometry of feather dimensions across bird taxa. Allometric relationships are commonly described by power-law equations of the form:

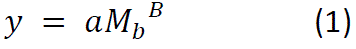

where scaling exponents (*B*) represent how traits (*y*) change in proportion to changes in body mass (*M_b_*) (Peters 1983). This scaling can manifest as either isometric scaling, wherein traits grow in direct proportion to body size maintaining geometric similarity of the traits (Alexander et al. 1979), or allometric scaling, which reflects a disproportionate change in a trait relative to body size (Huxley 1932). Allometric relationships can either be positively or negatively allometric. The particular scaling exponents expected in each type of relationship depend on the dimensionality of the trait in question. Mass is proportional to volume, with dimensions of length^3^. The *B* for similar volumetric traits under a condition of isometry is expected to be 1, while *B* < 1 or *B* > 1 would indicate negative or positive allometry, respectively. In cases where the dimensions of the trait are mismatched with respect to mass (e.g, if the trait is a length, with dimensions of length to the first power or an area, with dimensions of length squared), the isometric expectations would be *B*=0.33 and *B*=0.67, respectively.

Scaling relationships can have profound implications for the structure and function of particular traits. For example, in birds, because wing size (area) scales isometrically (*M* ^0.67^), birds maintain geometric similarity of their wings across body size (Taylor and Thomas 2014; but see Rayner 1988). The functional implication of this isometry is that large birds are at a comparative disadvantage because the scaling of their wings leads to relatively less wing surface area to produce lift during flight, leading to generally higher wing loading in larger birds (Warham 1977; Vogel 1981). A key component of the allometric scaling in wings lies in the morphology of the primary flight feathers, which themselves show some distinct patterns of allometry with respect to the bird’s size (Worcester 1996). Prior studies of allometric scaling in flight feathers found isometric scaling, but did not study the entire series of primaries (Worcester 1996). As a result, the scaling relationships across the series of flight feathers with respect to body size remains largely unknown. In this study, we measured the scaling relationships of the eleven distal-most flight feathers with three complementary predictions. First, since bird wings are known to scale isometrically (Taylor and Thomas 2014), we predicted similar isometric scaling of feather lengths. Second, if allometry were to exist among the lengths of the feathers, we predicted that the strongest allometry should be near the wingtip, as predicted by the aerodynamic gradient along the wing. Finally, if modularity existed within the wing, we expected to see stepwise changes in the scaling relationship along the wing, corresponding with the modules.

To address these open questions regarding the evolution of the flight feathers, we measured the lengths of the distal eleven flight feathers in a taxonomically broad sample of preserved wings and tested for morphological modularity among the flight feathers. Our sample included the entire series of primaries, and in some taxa, a few distal secondaries. We used two metrics to evaluate whether the evolutionary tempo of the feathers follows the predicted gradient of mechanical sensitivity: morphological disparity (a phylogenetically-informed measurement of shape variance among species) and *σ^2^*(the evolutionary rate parameter). We also measured the scaling relationships of each feather with respect to body mass to assess whether feathers scale isometrically or allometrically. Finally, we asked whether any morphological modularity exists within the distal flight feathers. Our results indicate that the pace of morphological evolution, particularly of the lengths of the flight feathers, follows a gradient pattern similar to that of the aerodynamic and inertial gradients generated by flapping flight. We also find that the majority of the feather sequence shows a pattern of isometric scaling, with the exception of the two most distal feathers, which scale with negative allometry. We conclude that the physical gradient generated by flapping flight is the underlying process driving feather evolution within the wings.

## Methods

Using digital calipers, we measured the lengths (± 0.1mm) of the 11 distal flight feathers from 509 wing specimens from 214 species in the collections at the North Carolina Museum of Natural Sciences (see Figure 1C, supplementary figure S1). This captured the full series of primary feathers and one or two of the distal-most secondary feathers (depending on the species). The relative lengths of these feathers determine the geometry of the wingtip (Fig. 1). We also recorded body mass from the museum tag data, when available (332 of 509 specimens). Some species in our sample did not have body mass (*M_b_*) recorded on their tags (49 of 214 species). For these samples, we used the mean values in the CRC Handbook of Avian Masses (Dunning Jr. 2007). Body size might exert a large influence on the subsequent analyses (Klingenberg 2016), so we used two approaches to nondimensionalize our feather measurements: 1) we divided the feather lengths by *M* ^1/3^ to transform the feather lengths into mass-specific length, and 2) we regressed the species-mean feather lengths against species-mean body mass and used the residuals as a measure of body size-independent feather length (‘*pgls*’ function, *caper* package). We tested the robustness of these methods by calculating Pearson’s Correlation Coefficient between the mass-transformed feather lengths and the residuals of the feather length – body mass relationship as the response (function ‘*cor.test*’, method= “Pearson”, *stats* package). Because these methods produced highly similar results (all *r* > 0.94, see Supplementary Figure S2), and because the mass-transformed lengths are more readily interpretable, we used those for subsequent analyses. All analyses were conducted in the R Statistical Computing Environment version 4.1.0 (R Development Core Team 2022).

Phylogenetically-explicit analyses were based upon the Jetz. *et al*. (2012) supertree (Ericson backbone; Figure 1C) from Birdtree.org (Rubolini et al. 2015), which includes 10,000 Bayesian posterior draws of the tree. We pruned the tree to include only the 214 species in our feather dataset using the ‘*drop.tip’* function (library “*ape*”, Paradis et al. 2004). To ensure the robustness of our analyses to different phylogenetic trees, we compared results using the Jetz. et al phylogeny (2012) with those obtained under the McTavish et al. tree (2024).

## Allometric scaling of flight feathers

We used phylogenetic generalized least squares (PGLS) regression to study the relationships between the lengths of each flight feather in the series and body mass. For each of the 11 measured feathers, we constructed a log-log scale PGLS model with species-mean feather length (log_10_(length)) as the response variable and body mass (log_10_(*M*_b_)) as the predictor using the *pgls* function in the *caper* package in R (Orme et al. 2018). We used a maximum likelihood optimization of Pagel’s *λ* (Pagel 1999), with specified bounds between 0.0001 and 1. We repeated these models using mass-specific measures (mm/g) of feather length as the response. To assess whether each feather showed significant allometry, we compared the measured slope to the predicted slope under a condition of isometry (0.33 for linear measures such as feather length) using a two-tailed t-test of the scaling model for each of the feathers in the series.

## Phylogenetic signal and evolutionary conservatism

We studied whether the lengths of of the flight feathers (mostly primaries, but also distal secondaries in some taxa) show evidence of evolutionary conservatism (more similarity than expected by the phylogenetic relationships between species; Harvey and Pagel 1991) by measuring the phylogenetic signal of each of the eleven measured feathers. We used the pruned Jetz et al. (2012) supertree which encompassed 214 branches. The tree we used was deposited at opentree.org (Accession number: TBD). We measured phylogenetic signal among the dimensional feather measurements using two complementary metrics: Blomberg’s *K* (Blomberg et al. 2003), and Pagel’s *λ* (Pagel 1999). We implemented these tests using the ‘*phylosig’* function in the *phytools* package (Revell 2012). Finally, we also calculated *K* and *λ* for each feather using the McTavish tree, and compared the Jetz and McTavish results using Pearson’s correlation coefficients (‘cor.test’ function, method=”pearson”).

Blomberg’s *K* (Blomberg et al. 2003) quantifies the similarity among related species relative to expectations under a Brownian motion model of evolution along branches of equal length. *K* assesses whether trait values change randomly in both direction and magnitude across the phylogeny and ranges from 0 to ∞. A value of *K* = 1 indicates that observed phenotypic values match those predicted by Brownian motion. When *K* < 1, related species are less similar than expected, and is considered evidence against phylogenetic niche conservatism (Losos 2008; Wiens et al. 2010; Cooper et al. 2011). Conversely, *K* > 1 indicates greater similarity than expected, potentially implying evolutionary conservatism, though this alone does not confirm it (see Wiens et al. 2010). To assess significance, we compared observed *K* values to those generated by iterative tip-shuffling randomization (Kembel et al. 2010), which removes phylogenetic structure to create a null expectation. A measured *K* exceeding this null suggests conservatism, while a lower value implies adaptive divergence.

Additionally, we estimated Pagel’s *λ* (Pagel 1999) with a maximum likelihood optimization (Revell 2012). *λ* represents the degree to which a clade’s phylogenetic history predicts trait distribution at the tree tips, ranging from 0 to 1. When *λ* = 0, the phylogeny resembles a star tree, with all tips equally distant from a common ancestor. A *λ* of 1 indicates a structure consistent with Brownian motion given the tree’s topology and branch lengths. We computed *λ* for each primary feather using the ‘*phylosig*’ function in the *Phytools* R Package (Revell 2012) with 1,000 simulations to test if the observed value differed from 0. We further assessed significance by performing likelihood ratio tests, comparing observed *λ* values to models where *λ* = 0 (no phylogenetic signal) and *λ* = 1 (Brownian evolution).

Blomberg’s *K* and Pagel’s *λ* capture distinct but complementary aspects of phylogenetic signal. *K* represents the ratio of among-species variance to the variance of independent contrasts (Blomberg et al. 2003), whereas *λ* describes the scale of correlation in trait values across the phylogeny (Pagel 1999). Both indexes converge at 1.0, indicating evolution consistent with Brownian motion. As *λ* approaches 0, phylogenetic relatedness becomes a weaker predictor of trait variation, while *K* clarifies whether trait divergence or conservation deviates from the Brownian expectation.

## Evolution of the flight feathers: rate, disparity and modularity

We calculated two metrics to describe the morphological divergence in the flight feathers: evolutionary rate and morphological disparity. The sigma-squared (*σ^2^*) evolutionary rate parameter quantifies the variance in trait evolution per unit time under a Brownian motion model, reflecting the rate at which traits diverge among lineages (Felsenstein 1985). To do this, we fit five different models of phenotypic evolution (Brownian motion, BM; Ornstein-Uhlenbeck, OU; early burst, EB; rate trend, RT; and white noise, WN) to the feather data using the ‘*fitContinous*’ function in the *geiger* R package (Harmon et al. 2008; Pennell et al. 2014). We used an Akaike’s Information Criterion with small sample sizes correction (AICc) to determine the most suitable model for each feather in the primary series and estimated *σ^2^* using that model. Morphological disparity is the extent of variation among taxa in a morphological trait (Stanley 1985; Gould 1991; Guillerme et al. 2020). We compared morphological disparity in each of the flight feathers among bird taxa using the ‘*dispRity’* function in the *dispRity* package (Guillerme 2018) in R. These analyses yielded a measurement of evolutionary rate and morphological disparity, across our taxonomic sample, for each position along the flight feather series. To assess whether *σ^2^* and morphological disparity were higher near the distal end of the flight feather series (i.e., at the tip of the wing), we used Generalized Least Squares (GLS) regression models with the estimates of *σ^2^* under each of the evolutionary regimes (i.e., BM, OU, EB, RT and WN) as the dependent variables and feather position (1 through 11) as the predictor. We conducted a similar GLS model with morphological disparity treated as the dependent variable in each model and feather position as the predictor. We used the *‘gls’* function in the *nlme* package (Pinheiro et al. 2013) for these regressions. We then repeated these models and included terms that account for spatial autocorrelation within the feather series, and used likelihood ratio tests to determine whether the models that included the autocorrelation terms were favored over the simple GLS models. We compared the evolutionary rate estimates from the Jetz phylogeny to those produced with the McTavish phylogeny by calculating the Pearson’s correlation coefficient for each evolutionary model (‘cor.test’ function, method=”pearson”).

Finally, we used the covariance ratio (*CR*) test from Adams (2016) to assess whether there is any modular organization among the primary feathers. When modularity exists among a suite of morphological traits, those traits that compose any given module are expected to covary more strongly with each other than they do with traits outside of the module, or within other modules (Klingenberg 2008; Adams 2016; Zelditch and Goswami 2021). The CR test compares covariance among traits within a hypothetical module to covariance among the modules. The test statistic (*CR*) ranges from 0 to positive infinity. Values between 0 and 1 indicate greater covariance within proposed modules than among them, which in turn signals morphological modularity. *CR* greater than 1 indicates a lack of modularity (morphological integration; Adams 2016). We implemented the *CR* test using code provided in the supplement of Adams’ description of the method (2016). To test for modularity, we binned the feathers (across taxa) into groups of two, three, and four contiguous feathers and tested for modularity among the bins (see Figure 4 and Figure S3). We iterated our tests on both the fully dimensional measurements of feather lengths and also on the dimensionless relative feather lengths in case feather size influenced the outcome of the CR tests.

## Results

Average feather length ranged from 19.3 mm (a hummingbird, genus *Selasphorus*) to 403 mm (the secretary bird, *Sagittarius serpentarius*). Because the lengths of the feathers vary across several orders of magnitude, we scaled their lengths relative to body mass (*M* ^1/3^). This non-dimensionalized the feather lengths and removed body size bias from our analyses (Adams 2013). The relative feather length ranged from 6.4 to 46.4. The longest feather in the sequence varied among species, but in general, relative feather length increased toward the tip of the wing (see Figure 2A).

**Figure 2.**
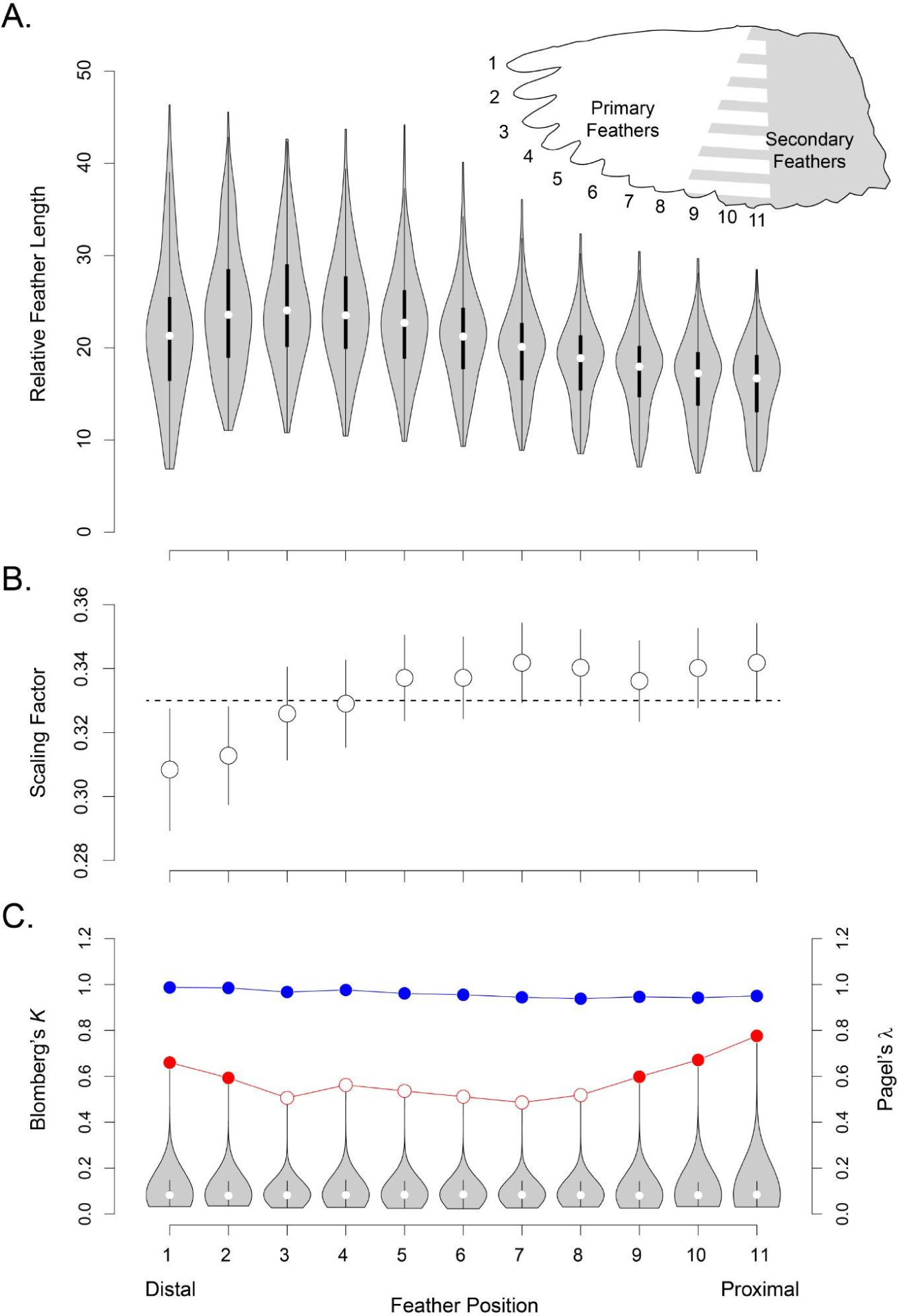
Relative lengths and phylogenetic signal of the primary flight feathers. A) Boxplot depicting the lengths of each feather in the primary sequence. Feather lengths were non-dimensionalized by dividing them by M_b_^0.33^. B) Scaling factor was determined by fitting a phylogenetic generalized least-squares model to the log-log relationship between each feather in the series and body mass. Points denote the slope coefficient from the PGLS models, which correspond with the scaling factor, and the error bars show the standard error associated with the slope estimation. The open circles denote feathers that did not deviate significantly from isometry (M_b_^0.33^, shown by the dashed line). C) Phylogenetic signal was measured with both Blomberg’s *K* (red) and Pagel’s *λ* (blue). Values that differ significantly from 1.0 are shown as open circles; those that are non-distinct from 1.0 are closed circles.

## Allometric scaling of flight feathers

First, we used phylogenetic generalized least-squares regression to describe the scaling relationship between the lengths of each flight feather along the wing and body mass (*M*_b_), across taxa. We found that all feather lengths scaled isometrically with respect to body mass (approximately *M* ^0.33^), matching our prediction based on the known scaling of whole-wing area (Taylor and Thomas 2014). This was true when we pooled all of the primary feathers (overall scaling relationship: *M* ^0.34^), and also when we looked at each feather in the series individually (Figure 2B and Table 1).

**Table 1.** Scaling relationships of the flight feathers.

| Feather | Scaling Factor ± SE | *P* |
| --- | --- | --- |
| p1 | 0.308 ± 0.019 | 0.194 |
| p2 | 0.313 ± 0.015 | 0.183 |
| p3 | 0.326 ± 0.015 | 0.613 |
| p4 | 0.329 ± 0.014 | 0.754 |
| p5 | 0.337 ± 0.013 | 0.779 |
| p6 | 0.337 ± 0.013 | 0.768 |
| p7 | 0.342 ± 0.012 | 0.496 |
| p8 | 0.340 ± 0.012 | 0.560 |
| p9 | 0.336 ± 0.013 | 0.825 |
| p10 | 0.340 ± 0.012 | 0.581 |
| s1 | 0.342 ± 0.012 | 0.492 |

## Phylogenetic signal and evolutionary conservatism

Next, we measured the phylogenetic signal among taxa for each of the primary feathers. We used two complementary indexes of phylogenetic signal: Pagel’s *λ* (Pagel 1999) and Blomberg’s *K* (Blomberg et al. 2003). Both metrics were significantly lower than 1 for the second through fourth primary feathers (Figure 2C, Table 2 and Supplementary Table S1). The phylogenetic signal results between the Jetz and McTavish trees were not correlated (*r* < 0.1 and *p* > 0.8 for both signal metrics), but they were qualitatively similar (Figure S3). Taken together, these results suggest that the distal feathers (excluding the first primary) have evolved faster than expected under the neutral model of Brownian motion.

**Table 2.** Non-dimensional phylogenetic signal results.

| Feather | *K* | $\lambda$ | *p* (*K*=0) | *p* (*K*=1) | *p* ($\lambda$=0) | *p* ($\lambda$=1) |
| --- | --- | --- | --- | --- | --- | --- |
| p1 | 0.66 | 0.987 | 0.001 | 0.17 | 0 | 0 |
| p2 | 0.593 | 0.985 | 0.001 | 0.071 | 0 | 0 |
| p3 | 0.506 | 0.967 | 0.001 | 0.015 | 0 | 0 |
| p4 | 0.562 | 0.976 | 0.001 | 0.044 | 0 | 0 |
| p5 | 0.536 | 0.961 | 0.001 | 0.029 | 0 | 0 |
| p6 | 0.511 | 0.955 | 0.001 | 0.022 | 0 | 0 |
| p7 | 0.486 | 0.944 | 0.001 | 0.006 | 0 | 0 |
| p8 | 0.518 | 0.938 | 0.001 | 0.014 | 0 | 0 |
| p9 | 0.598 | 0.946 | 0.001 | 0.071 | 0 | 0 |
| p10 | 0.671 | 0.942 | 0.001 | 0.153 | 0 | 0 |
| s1 | 0.776 | 0.95 | 0.001 | 0.361 | 0 | 0 |

Unlike primaries 2-4, phylogenetic signal across the rest of the feather sequence was generally strong, with Pagel’s *λ* not statistically distinguishable from 1 (Figure 2C, Table 2 and Supplementary Table S1). Similarly, we found that *K* was also approximately 1.0 for most feathers (Figure 2C), indicating that the evolution of feather lengths along most of the wingspan (other than primaries 2-4) can be modeled by a Brownian motion process – that is, the evolution of primary feather length has proceeded in a random fashion and the distribution among taxa is explained primarily by their evolutionary relatedness. Consistent with the high phylogenetic signal, the best fitting evolutionary model for the feathers was a Brownian motion model of phenotypic evolution (supplementary Table 3). Taken as a whole, the phylogenetic signal results indicate evolutionary divergence in most primary feather lengths that is commensurate with the phylogenetic diversity among bird taxa, fitting neither a model of evolutionary conservatism nor of adaptive divergence, but with a strong signal of adaptive divergence near the tip of the wing.

## Evolution of the flight feathers: rate, disparity and modularity

Next, we studied whether evolutionary rate (*σ^2^*) and morphological disparity were greater at the distal tip of the wing. The estimated values of *σ^2^* also differed among the evolutionary models (Table 3). The OU model was the best fit in both the scaled and the non-scaled datasets (per AICw, see Table 3, Supplementary Table S2). We found decreasing trends of *σ^2^* across the primary feather series, from distal to proximal (i.e., that feathers closer to the wingtip evolved more rapidly than those closer to the body), for all four of the evolutionary models that we tested (see BM and OU results in Figure 3, full model results in Table 3). The alpha parameter (*ɑ*) in the OU models was also greatest near the wingtip (see supplementary table S3), perhaps indicating that the distal portion of the wing has the strongest selective regime. The non-evolutionary (white-noise) model yielded a similar pattern, though the estimated coefficients diverged strongly from the evolutionary models (Table 3, Supplementary Table S2). We also note that we found similarly-shaped trends (greater distally, decreasing more proximally) across all of the evolutionary models, regardless of the inclusion or not of a model term that accounts for autocorrelation among the feathers (Table 4). Finally, we found similar trends of *σ^2^* for both the Jetz and McTavish trees (Figure S3), suggesting these results are robust to differing phylogenetic hypotheses. Morphological disparity, essentially a measure of morphological divergence in feather dimensions among taxa, also followed a significant decreasing trend in the feather series (Table 3, Figure 2). The combination of increasing evolutionary rate and greater morphological disparity toward the wingtip suggest increasingly strong evolutionary divergence in feather length from the base to the tip of the wing, which is consistent with the phylogenetic signal results.

**Figure 3.**
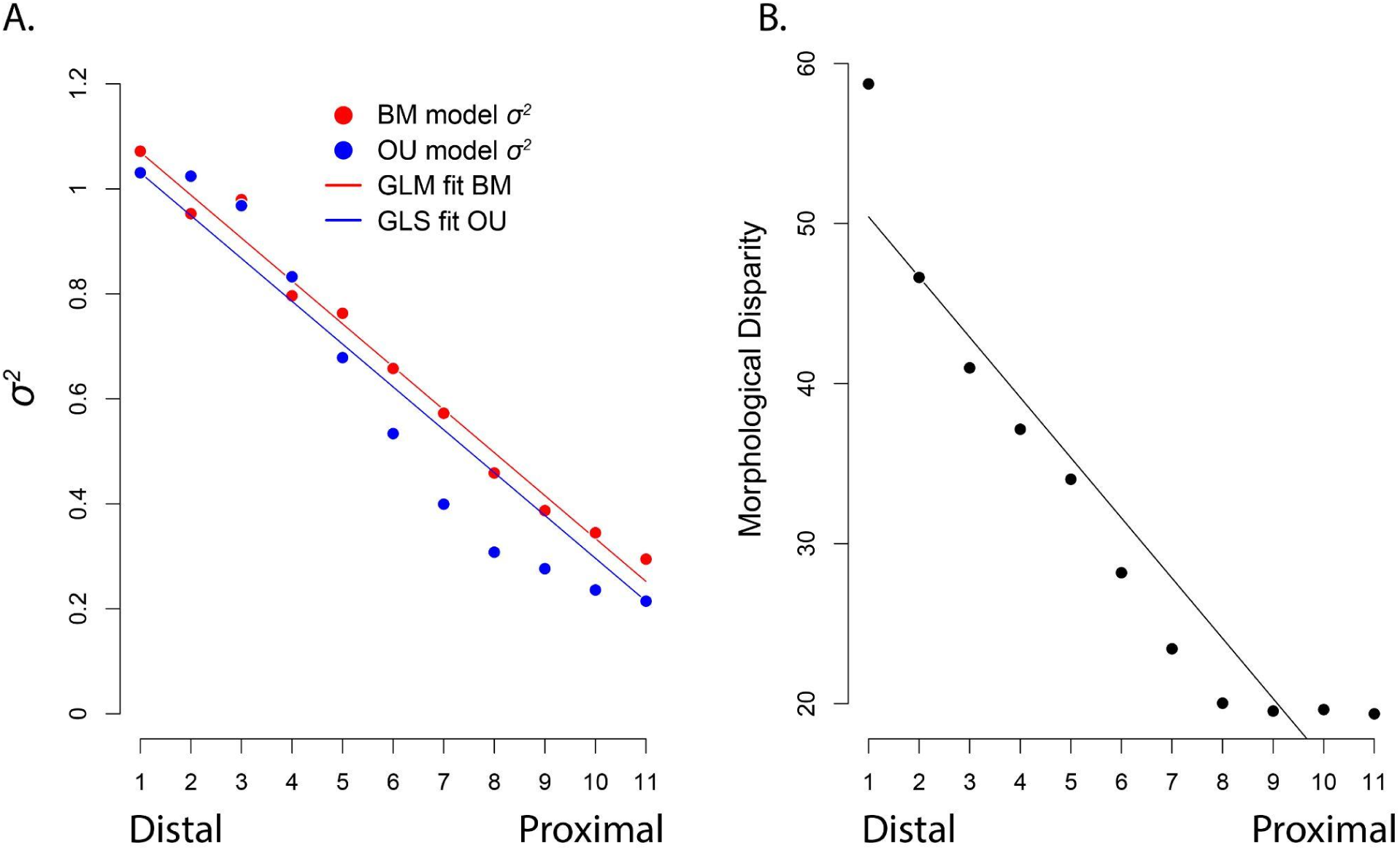
Evolutionary dynamics of the primary flight feathers. A) Evolutionary rate (*σ^2^*) measured via two models of evolution: Brownian motion (red) and Ornstein-Uhlenbeck (blue). We also measured *σ^2^* using a non-evolutionary white-noise model (see Table S1). B) Morphological disparity among taxa for the primary flight feathers. In both panes, points represent calculated values for each feather and the line depicts the results of a generalized linear fit to those values.

**Figure 4.**
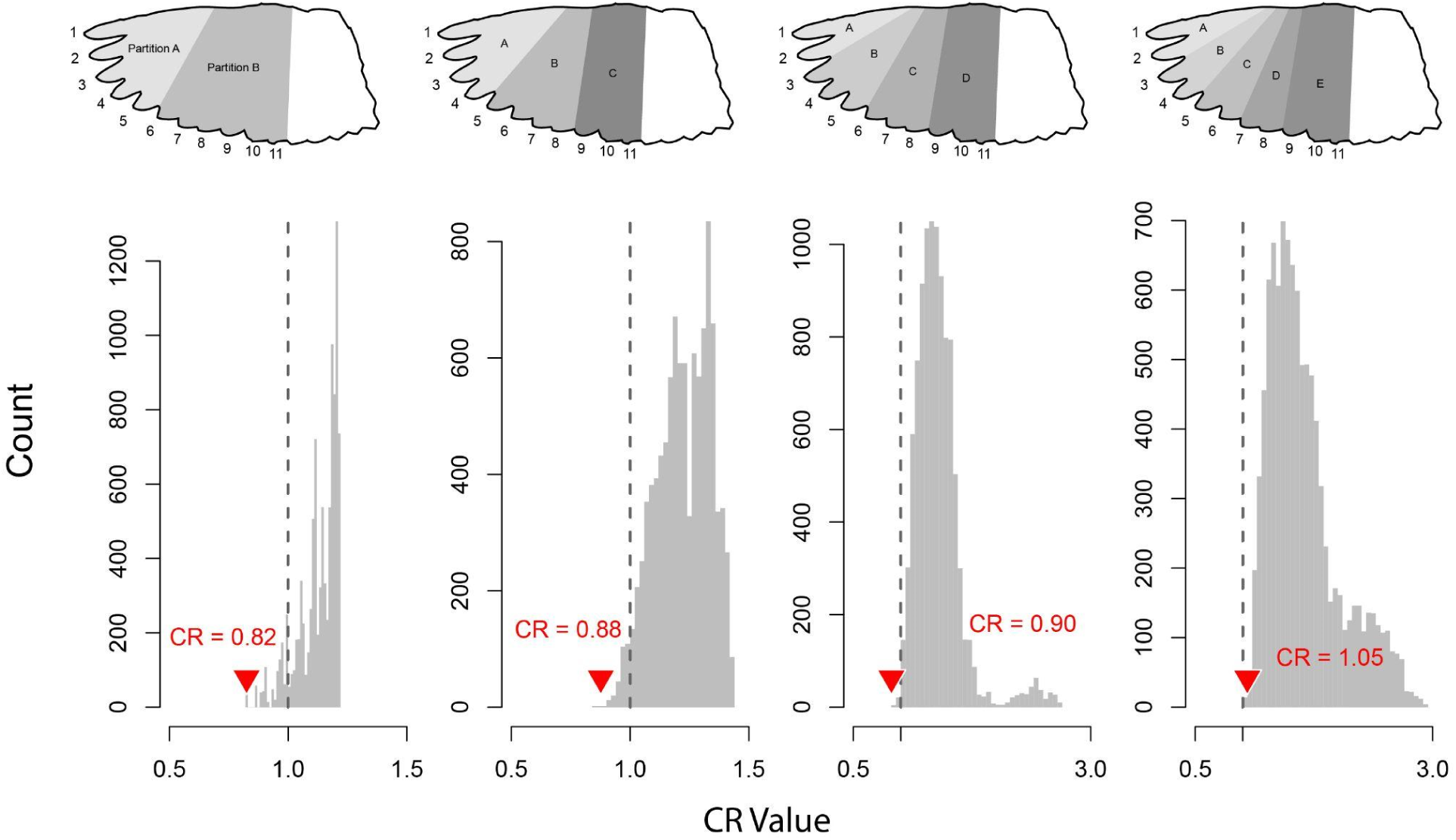
Modularity test of the non-dimensionalized primary feathers. Wing illustrations depict schemes of partitioning feathers into hypothetical modules. Feather lengths were non-dimensionalized by dividing them by M_b_^1/3^. Modularity was tested using the Covariance Ratio test from Adams (2016). CR < 1.0 indicates modularity, while CR > 1.0 indicates integrated evolution. The recovered CR values for each partitioning scheme are shown with an inverted red triangle. The gray background histograms show the result of a permutation test, randomly shuffling the feathers among the putative modules (*n* = 10,000).

**Table 3.** Results of evolutionary rate models of non-dimensional feather lengths. Evolutionary rate (**σ**^2^) estimated by each model (top line for each model) and AIC weights (bottom line for each model).

| Model | Feather1 | Feather2 | Feather3 | Feather4 | Feather5 | Feather6 | Feather7 | Feather8 | Feather9 | Feather10 | Feather11 |
| --- | --- | --- | --- | --- | --- | --- | --- | --- | --- | --- | --- |
| **BM** | 1.07 | 0.95 | 0.98 | 0.80 | 0.76 | 0.66 | 0.57 | 0.46 | 0.39 | 0.34 | 0.29 |
|  | 0.01 | 0.00 | 0.00 | 0.00 | 0.00 | 0.00 | 0.00 | 0.00 | 0.00 | 0.00 | 0.00 |
| **OU** | **1.03** | **1.02** | **0.97** | **0.83** | **0.68** | **0.53** | **0.40** | **0.31** | **0.28** | **0.24** | **0.21** |
|  | **0.91** | **0.92** | **1.00** | **1.00** | **1.00** | **1.00** | **1.00** | **1.00** | **1.00** | **1.00** | **1.00** |
| **EB** | 1.07 | 0.95 | 0.98 | 0.80 | 0.76 | 0.66 | 0.57 | 0.46 | 0.39 | 0.34 | 0.29 |
|  | 0.00 | 0.00 | 0.00 | 0.00 | 0.00 | 0.00 | 0.00 | 0.00 | 0.00 | 0.00 | 0.00 |
| **rate_trend** | 0.00 | 0.00 | 0.00 | 0.00 | 0.00 | 0.00 | 0.00 | 0.00 | 0.00 | 0.00 | 0.00 |
|  | 0.08 | 0.07 | 0.00 | 0.00 | 0.00 | 0.00 | 0.00 | 0.00 | 0.00 | 0.00 | 0.00 |
| **white** | 58.45 | 46.41 | 40.78 | 36.97 | 33.86 | 28.04 | 23.31 | 19.93 | 19.43 | 19.53 | 19.27 |
|  | 0.00 | 0.00 | 0.00 | 0.00 | 0.00 | 0.00 | 0.00 | 0.00 | 0.00 | 0.00 | 0.00 |
| **Disparity** | 58.72 | 46.63 | 40.97 | 37.14 | 34.01 | 28.17 | 23.42 | 20.02 | 19.53 | 19.62 | 19.36 |

**Table 4.** Evolutionary rate and disparity model coefficients.

| Model | Int_lm | Slope_lm | P_Coeff | Int_gls | Slope_gls | P_Coeff | LogLik_lm | LogLik_gls | LR | P_LRT |
| --- | --- | --- | --- | --- | --- | --- | --- | --- | --- | --- |
| $\sigma^2_{BM}$ | **1.150** | **-0.081** | <0.0001 \| <0.0001 | 1.152 | -0.082 | <0.0001 \| <0.0001 | 13.435 | 13.541 | 0.211 | 0.646 |
| $\sigma^2_{OU}$ | 1.171 | -0.097 | <0.0001 \| <0.0001 | **1.112** | **-0.082** | 0.9031 \| 0.0014 | 7.285 | 11.876 | 9.180 | 0.002 |
| $\sigma^2_{EB}$ | **1.151** | **-0.081** | <0.0001 \| <0.0001 | 1.152 | -0.082 | <0.0001 \| <0.0001 | 13.434 | 13.540 | 0.212 | 0.646 |
| $\sigma^2_{RT}$ | **0.000** | **0.000** | <0.0001 \| <0.0001 | 0.000 | 0.000 | <0.0001 \| <0.0001 | 94.330 | 94.341 | 0.021 | 0.884 |
| $\sigma^2_{WN}$ | 53.931 | -3.746 | <0.0001 \| <0.0001 | **62.368** | **-3.918** | 0.9364 \| 0.0071 | -29.745 | -25.398 | 8.693 | 0.003 |
| **Disparity** | 54.184 | -3.764 | <0.0001 \| <0.0001 | **62.661** | **-3.937** | 0.9364 \| 0.0071 | -29.787 | -25.441 | 8.693 | 0.003 |

We found conflicting evidence for modularity among the feather series. We tested for morphological modularity among the feathers under two regimes: 1) fully-dimensional, using the non-scaled lengths of the feathers, and 2) nondimensional, with feathers scaled relative to M ^0.33^. We were also concerned that body size might influence the modularity test because of the extreme difference in body size among our study taxa (2.7 g to 6,050 g). Among the flight feathers in the dimensionless dataset, the CR value was consistently less than 1.0 and significantly less than the null-expectation generated by the permutation test regardless of how we partitioned the feathers (see Figure 4, Table 5). In contrast, the CR values recovered for the non-scaled dimensional dataset were all greater than 1, suggesting morphological integration among the feathers. However, the calculated CR values were still significantly less than the null expectation from the permutation test (see supplemental Figure S3 and Table 5). We interpret this discrepancy as clear that the variation in body size influences the CR calculations. The CR method compares the variance within a putative module to the variance between modules, and in the case of the dimensional feather lengths, a large amount of variance within modules arises simply from body size. We have also shown that evolutionary dynamics (rate and disparity), and the lengths of the feathers all vary along the feather sequence in a gradient fashion, and it remains unclear whether such a gradient might cause the CR method to find modularity where none exists, and we therefore hesitate to interpret these results further.

**Table 5.** Morphological modularity results.

| Number of Modules | Dimensional CR | Non-dimensional CR | $p_{\text{dimensional}}$ | $P_{\text{non-dimensional}}$ |
| --- | --- | --- | --- | --- |
| 2 | 1.051 | 0.825 | 0.0015 | 0.0025 |
| 3 | 1.120 | 0.877 | 0.0006 | 0.0002 |
| 4 | 1.193 | 0.901 | 0.0002 | 0.0001 |
| 5 | 1.307 | 1.048 | 0.001 | 0.001 |

## Discussion

We measured the lengths of the primary flight feathers to study whether the evolution of feather length follows a root-to-tip gradient predicted by the aerodynamics of flapping flight (Weis-Fogh 1972). We further asked whether the flight feathers have evolved as an integrated morphological unit, or whether there is modularity within the feather sequence. We found that both evolutionary rate (*σ^2^*) and morphological disparity followed the gradient of forces experienced during flapping; morphological evolution of the feathers forming the wing surface follows a pattern predicted by flight physics. We also show that the lengths of the measured flight feathers scale isometrically with respect to body mass, confirming our prediction. Finally, we found conflicting and equivocal evidence for morphological modularity among the flight feather sequence, suggesting some spatial covariance in their morphology, but we did not find a clear delineation of the modules. We discuss each of these results in the following paragraphs.

The primary result of our study is that we show that the pace of evolution in the lengths of the flight feathers follows a gradient pattern along the length of the wing. Both evolutionary rate and morphological disparity increase from proximal to distal, tracking the gradient of aerodynamic force along the wing (Weis-Fogh 1972). A similar gradient pattern has been observed in the evolution of overall wing shape. Morphological disparity of four shape traits (chord, camber, cross-sectional thickness, and cross-sectional area) are greatest near the tip of the wing and decline toward its base (Rader and Hedrick 2023), following a gradient along the wing predicted by the distribution of aerodynamic forces that are experienced by flapping-winged fliers (Weis-Fogh 1972; Shyy et al. 2010). However, these earlier findings regarding the overall wing morphology did not account for the contributions of individual feathers to gradient patterns of evolution along the wing. Our results here indicate that the evolution of individual feather dimensions follows a similar pattern to the overall wing morphology.

The mechanical sensitivity of traits within biomechanical systems has been implicated as a mediator of evolutionary tempo and mode of those traits (Muñoz et al. 2017, 2018).

Specifically, traits that have the strongest influence over the functional output of biomechanical systems also tend to show signs of adaptive evolution (i.e., fitting a model of directed evolution such as Ornstein-Uhlenbeck) coupled with comparatively high rates of evolutionary divergence. 4-bar linkage systems, such as those found in the jaws of many bony fish or in the raptorial appendages of mantis shrimp, represent some of the most clear examples of the link between mechanical sensitivity and morphological evolution. In these systems, the length of the link dictates their contribution to the overall output of the system through direct kinematic transmission, and those links that have the greatest impact on output evolve fastest (Muñoz et al. 2018). By contrast, the influence of specific wing shape traits on the output of the biomechanical system (the overall wing) is less apparent. Flapping wings experience complex fluid-structure interactions where unsteady aerodynamic forces dynamically deform the flexible wing, creating nonlinear feedback between fluid flow and structural response (Lentink and Dickinson 2009; Shyy et al. 2010; Chin and Lentink 2016). This complexity makes it difficult to isolate the influence of wing shape traits, as their effects depend on interactions between vortex dynamics, material properties, and wing kinematics (Combes and Daniel 2003a,b; Zhu 2007; Shoele and Zhu 2013; Colognesi et al. 2021). Nonetheless, the concordance of patterns between aerodynamic forces and morphological evolution in the wing shown here suggests that the mechanical sensitivity framework is also useful for predicting the evolution of fluid-structure systems. Our results therefore contribute to the emergent body of evidence that suggests that the relationship between mechanical sensitivity and morphology is an important driver in the evolution of biomechanical systems.

A second set of results from our report is that the flight feathers scale isometrically across the range of body size in birds. Mean primary feather length scaled isometrically (M ^0.34^) when we pooled the feathers in our sample, and we found similar results when we analyzed each individual feather position across taxa. These results confirmed our prediction, which was based on a priori knowledge of the scaling of whole wing area (Taylor and Thomas 2014), and they are also in agreement with prior studies of feather allometry (Worcester 1996; Wang et al. 2012; Sullivan et al. 2019). Collectively, our results suggest that the lengths of the distal flight feathers are evolutionarily conserved, at least when compared to other aspects of avian morphology such as bill shape (Cooney et al. 2017), the talons of raptors (Fowler et al. 2009; Tsang et al. 2019), and the range of motion in the wing joints (Baliga et al. 2019). The conserved nature of the flight feathers is in concordance with other static aspects of wing morphology including wing length, chord, aspect ratio, wing area, and the lengths of the wing bones (Taylor and Thomas 2014; Baliga et al. 2019).

Other aspects of flight feathers scale allometrically, with their biological nature influencing not only their own size but also the overall wing surface. For instance, feather stiffness scales with negative allometry, decreasing significantly as body size increases despite maintaining geometric similarity (Worcester 1996). This reduction in structural strength at larger sizes may impose an upper limit on feather dimensions and, consequently, on the planform area of the handwing. Additionally, feather keratin is metabolically inert, meaning feathers degrade over time without repair and must be periodically molted and regrown. This process affects multiple aspects of avian biology (Miller 1941; Tucker 1991; Murphy 1996; Chandler et al. 2010). Since feather production is energetically costly (Murphy 1996; Guillemette et al. 2007), these constraints may further limit feather size. Moreover, molt duration increases disproportionately in larger birds (Rohwer et al. 2009; Jenni and Winkler 2020; Jenni et al. 2020), potentially restricting the overall scaling of wing size relative to body size.

The isometric scaling of wings and of their feathers presents a mechanical paradox because of the dimensional mismatch between wing area and body mass. Lift forces are, all else being equal, a product of wing area. Body mass on the other hand is proportional to volume, and this difference means that the ratio of body mass to wing area, or wing loading, increases disproportionately across avian body size (Warham 1977; Vogel 1981). This paradox might emerge from variable evolutionary lability between the components of the wing and body size. Whereas selection seems to generally favor large body size (Kingsolver and Pfennig 2004, 2007), the length of feathers may be constrained by the physical stiffness of their keratin (Worcester 1996; Wang et al. 2012). The positive allometry of wing loading presents several disadvantages, particularly for takeoff, slow flight, and maneuverability. Higher wing loading requires faster flight speeds (Alerstam et al. 2007) or greater muscular power output (Tobalske and Dial 2000), which can increase energy expenditure and reduce flight efficiency. Additionally, birds with high wing loading face greater difficulty in generating sufficient lift during takeoff, often requiring long running starts or specialized launch behaviors, as seen in large waterfowl and terrestrial birds. Evolutionary resolutions to the wing loading paradox may therefore emerge in physiological or behavioral traits that have greater evolutionary lability (Blomberg et al. 2003) than feathers display, or there may be other aspects of wing shape (particularly 3-dimensional attributes like camber) that contribute to enhancing lift production for a given wing area (Brown 2001; Waldrop et al. 2020).

Our results also have implications for the understanding of modular structures. Bird wings are composed of at least two morphological modules, the handwing and the armwing (Rader and Hedrick 2023; Orkney and Hedrick 2024). Here, we demonstrate that the primary feather sequence also shows evidence for morphological modularity, but evidence for the particular partitioning of feathers is equivocal. The flight feathers are serially homologous anatomical elements along the length of the wing, and the base-to-tip gradient of aerodynamic forces exposes them to a gradient of evolutionary pressures. Other examples of serially homologous anatomical structures superimposed over regionalized function show strong morphological modularity. For example, the mammalian backbone is organized into repeating vertebrae that are regionalized into cervical, thoracic and lumbar modules, each with its own evolutionary dynamics (Jones et al. 2018). We did not find similar clarity in the regionalization of morphology among avian flight feathers, perhaps because the regionalization of function is less discrete. In the mammalian backbone, the thoracic region is associated with the ribcage and the shoulder girdle, while the lumbar region is associated with the abdominal region and the cervical region with support for the cranium. By contrast, the feathers have no such strong delineations, and are instead superimposed across a square-powered force gradient. The appearance of modularity among the feathers may be an emergent property of the exponential nature of the gradient, a phenomenon which might merit explicit study.

Our finding that the most significant evolutionary divergence in feather dimensions is near the wingtip is also consistent with the growing body of literature demonstrating numerous links between handwing shape and other aspects of avian biology such as habitat preference or flight behavior (e.g., Claramunt et al. 2012; Taylor and Thomas 2014; Stoddard et al. 2017; Sheard et al. 2019, 2020). Wingtip shape influences factors such as lift-to-drag ratio, maneuverability, and energy efficiency during flight (Norberg 1990; Pennycuick 2008). Species adapted for high-speed or long-distance flight, such as swifts and albatrosses, often have elongated, pointed wingtips that reduce induced drag and enhance aerodynamic efficiency (Vogel 1981; Rayner 1988; Taylor and Thomas 2014). In contrast, birds that rely on rapid takeoff, agile maneuvering, or flight in cluttered environments, such as forest-dwelling passerines or raptors, tend to have more rounded or slotted wingtips that improve lift production and control at low speeds (Savile 1957; Rayner 1988; Tucker 1993; Nudds et al. 2004; KleinHeerenbrink Marco et al. 2017). Beyond flight performance, wingtip morphology may also reflect ecological specializations (Rayner 1988; Lockwood et al. 1998), predator-prey interactions (Swaddle and Lockwood 1998), and evolutionary trade-offs between flight efficiency and other life-history traits (Claramunt et al. 2012; Stoddard et al. 2017; Derryberry et al. 2018; Sheard et al. 2019, 2020). The adaptation to different habitats, life histories, and flight behaviors is reflected in the enhanced evolutionary rates that we find at the tip of the wing.

We also note that there are some caveats and limitations to our study. Our interpretations are restricted to the primary feathers and one or two distal secondaries in some taxa. We are thus unable to comment on evolutionary variation among the feathers that comprise the rest of the aerodynamic surface of the wing (the full series of secondaries, tertials, and coverts). There is significant variation in the number of secondary feathers in the wing, with proportionally long wings possessing a greater number of secondary feathers (Deeming et al. 2024). Aspect ratio (the ratio of wing length over the mean width of the wing) has repeatedly been shown to be one of the primary axes of morphological variation among bird wings (Vogel 1981; Rayner 1988; Nudds et al. 2004; see Taylor and Thomas 2014; Rader et al. 2020), and has been linked to several aspects of gliding flight performance (Waldrop et al. 2020; Ding 2024). As such, an explicit examination of the evolutionary divergence in secondary feather count and dimensions may be useful to describe the evolutionary transition from shorter wings in taxa that rely heavily on flapping to the longer wings that are characteristic of soaring birds. Finally, wing shape is influenced by not only the lengths of the feathers, which comprise the aerodynamic surface of the wing, but it is also a dynamic property that varies with wing posture in both biomechanically and evolutionarily relevant ways (Lentink et al. 2007; Baliga et al. 2019; Harvey et al. 2019, 2022). How the relative lengths of the flight feathers interact with mobility of the limb joints to determine the morphological envelope of the handwing might be a fruitful avenue for further study.

## Author contributions

Conceptualization: JAR

Methodology: JAR

Investigation: JAR, EBB, MEP

Visualization: JAR

Supervision: DRM

Writing—revised draft: JAR, EBB, MEP, DRM

Writing—review & editing: JAR, DRM

## Acknowledgements

We thank the North Carolina Museum of Natural Sciences, and especially Brian O’Shea and John Gerwin, for access to specimens. Patrick W. Kelly and Sean A. S. Anderson provided valuable feedback on this manuscript. Finally, we thank Tyson L. Hedrick for his support throughout this project.

## Data Accessibility Statement

Feather measurements and species-level summary will be archived with Figshape, the phylogenetic trees in OpenTree, and analysis scripts in Zenodo and will be made available upon manuscript acceptance.

---

**Table S1.** Dimensional phylogenetic signal results.

| Feather | *K* | $\lambda$ | *p* (*K*=0) | *p* (*K*=1) | *p* ($\lambda$=0) | *p* ($\lambda$=1) |
| --- | --- | --- | --- | --- | --- | --- |
| p1 | 0.836 | 0.991 | 0.001 | 0.555 | 0 | 0.079 |
| p2 | 0.8 | 0.991 | 0.001 | 0.432 | 0 | 0.042 |
| p3 | 0.767 | 0.987 | 0.001 | 0.364 | 0 | 0.023 |
| p4 | 0.777 | 0.986 | 0.001 | 0.391 | 0 | 0.017 |
| p5 | 0.73 | 0.979 | 0.001 | 0.281 | 0 | 0.001 |
| p6 | 0.763 | 0.986 | 0.001 | 0.384 | 0 | 0.006 |
| p7 | 0.74 | 0.982 | 0.001 | 0.284 | 0 | 0.001 |
| p8 | 0.715 | 0.98 | 0.001 | 0.258 | 0 | 0 |
| p9 | 0.694 | 0.98 | 0.001 | 0.21 | 0 | 0.001 |
| p10 | 0.727 | 0.979 | 0.001 | 0.278 | 0 | 0 |
| s1 | 0.743 | 0.979 | 0.001 | 0.32 | 0 | 0 |

**Table S2.** Results of evolutionary rate models of dimensional feather lengths. Evolutionary rate (**σ**^2^) estimated by each model (top line for each model) and AIC weights (bottom line for each model).

| Model | Feather1 | Feather2 | Feather3 | Feather4 | Feather5 | Feather_6 | Feather7 | Feather8 | Feather9 | Feather10 | Feather11 |
| --- | --- | --- | --- | --- | --- | --- | --- | --- | --- | --- | --- |
| BM | **97.02** | 114.30 | 125.79 | 123.67 | 125.59 | 104.82 | 96.22 | 85.87 | 80.01 | 73.33 | 64.68 |
|  | **0.38** | 0.27 | 0.20 | 0.14 | 0.01 | 0.05 | 0.00 | 0.00 | 0.02 | 0.00 | 0.00 |
| OU | 84.78 | **99.12** | **104.69** | **101.48** | **94.96** | **84.12** | **73.24** | **67.73** | **65.67** | **55.89** | **48.80** |
|  | 0.32 | **0.48** | **0.58** | **0.67** | **0.96** | **0.88** | **0.99** | **0.99** | **0.93** | **0.98** | **0.99** |
| EB | 97.03 | 114.31 | 125.80 | 123.68 | 125.60 | 104.83 | 96.23 | 85.87 | 80.01 | 73.34 | 64.68 |
|  | 0.14 | 0.10 | 0.07 | 0.05 | 0.00 | 0.02 | 0.00 | 0.00 | 0.01 | 0.00 | 0.00 |
| rate_trend | 74.30 | 67.41 | 58.22 | 51.45 | 31.91 | 35.26 | 24.01 | 16.40 | 19.36 | 17.90 | 18.60 |
|  | 0.16 | 0.15 | 0.15 | 0.13 | 0.02 | 0.05 | 0.01 | 0.01 | 0.04 | 0.01 | 0.01 |
| white | 6371.60 | 6928.02 | 7224.07 | 7096.21 | 6693.64 | 5693.05 | 4940.13 | 4293.24 | 3989.06 | 3651.08 | 3289.24 |
|  | 0.00 | 0.00 | 0.00 | 0.00 | 0.00 | 0.00 | 0.00 | 0.00 | 0.00 | 0.00 | 0.00 |
| Disparity | 6401.52 | 6960.55 | 7257.98 | 7129.52 | 6725.06 | 5719.77 | 4963.32 | 4313.40 | 4007.79 | 3668.22 | 3304.69 |

**Table S3.** Alpha parameter (*ɑ*) for Ornstein-Uhlenbeck models of feather evolution.

| Feather | p1 | p2 | p3 | p4 | p5 | p6 | p7 | p8 | p9 | p10 | s1 |
| --- | --- | --- | --- | --- | --- | --- | --- | --- | --- | --- | --- |
| $\alpha$ | 0.0047 | 0.0087 | 0.0111 | 0.0108 | 0.0093 | 0.0091 | 0.0079 | 0.0075 | 0.0055 | 0.0041 | 0.0032 |

**Figure S1.**
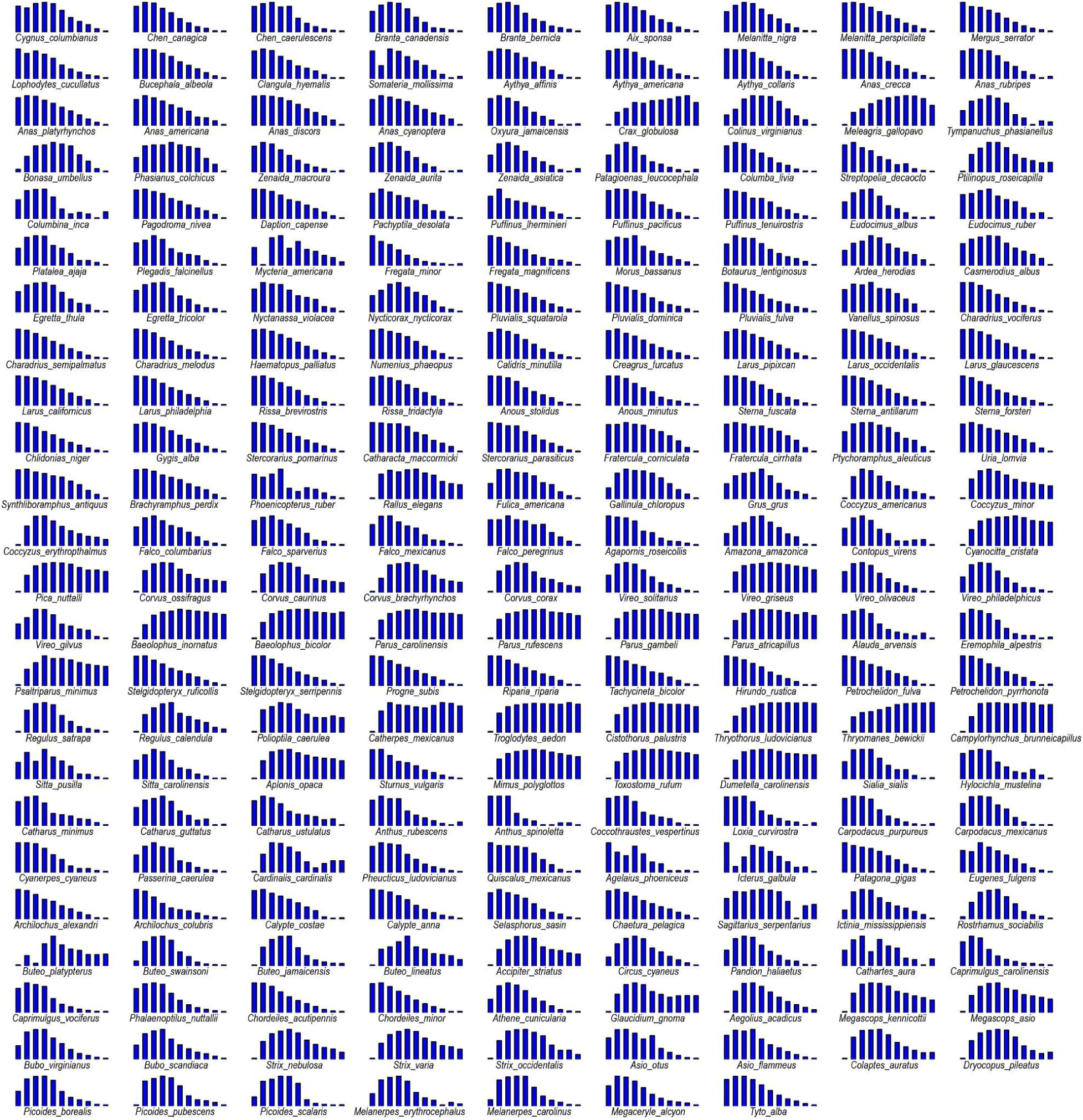
Relative lengths of feathers for the 214 species included in this study. Subplots are similar to Panel B in Figure 1. Species are arranged in order of the tree tips (Tip 1 is on the bottom of the tree, Tip 214 is the top).

**Figure S2.**
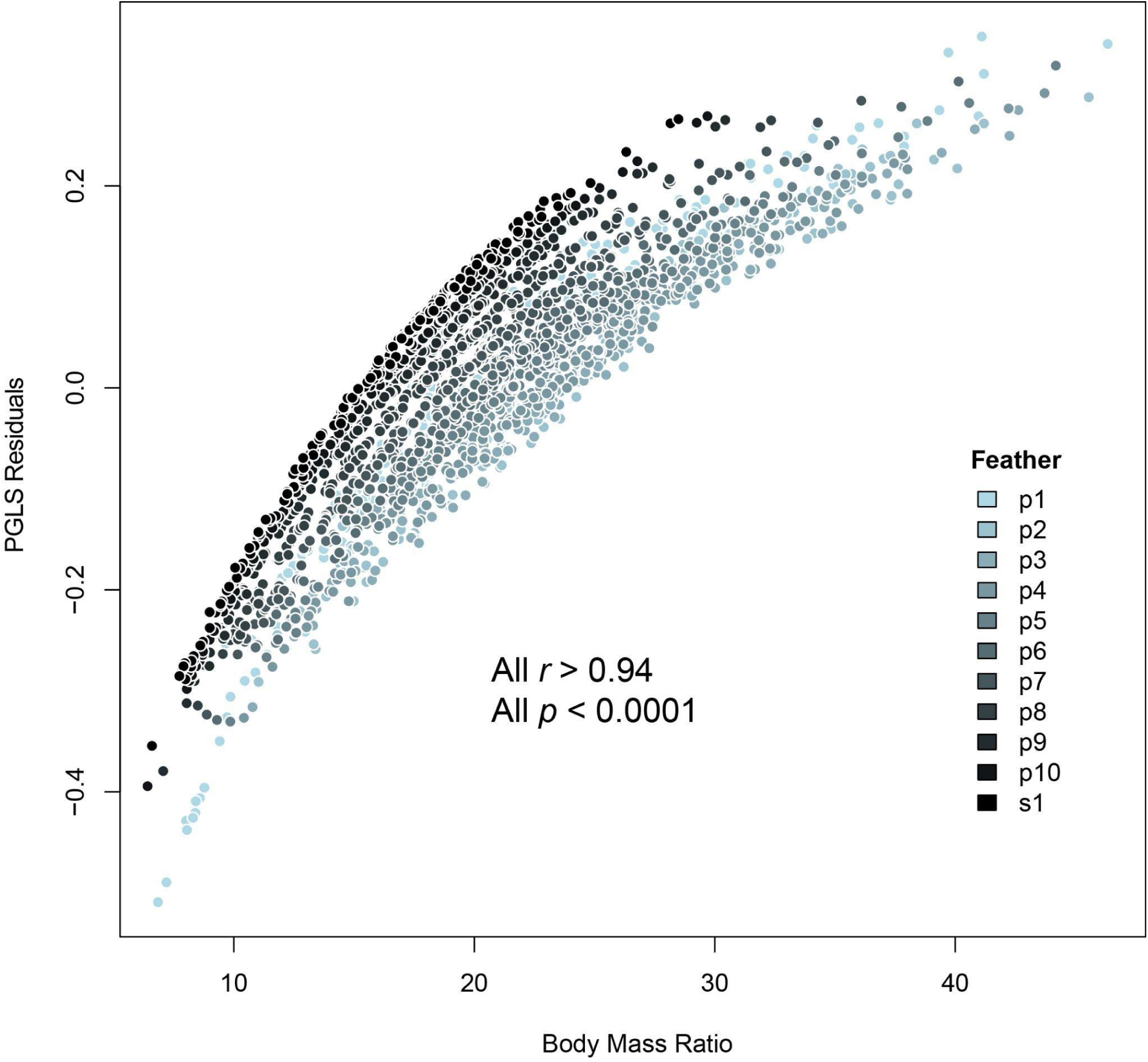
Comparison of feather non-dimensionalization methods. We tested two methods for non-dimensionalizing feather lengths with respect to body mass (M_b_): 1) a ratio of feather length over Mb, and 2) residuals of log-log PGLS regressions with feather length as the response and Mb as predictor. These methods produced results that were highly correlated (Pearson’s Correlation Coefficient, *‘cor.test’* function, *stats* package).

**Figure S3.**
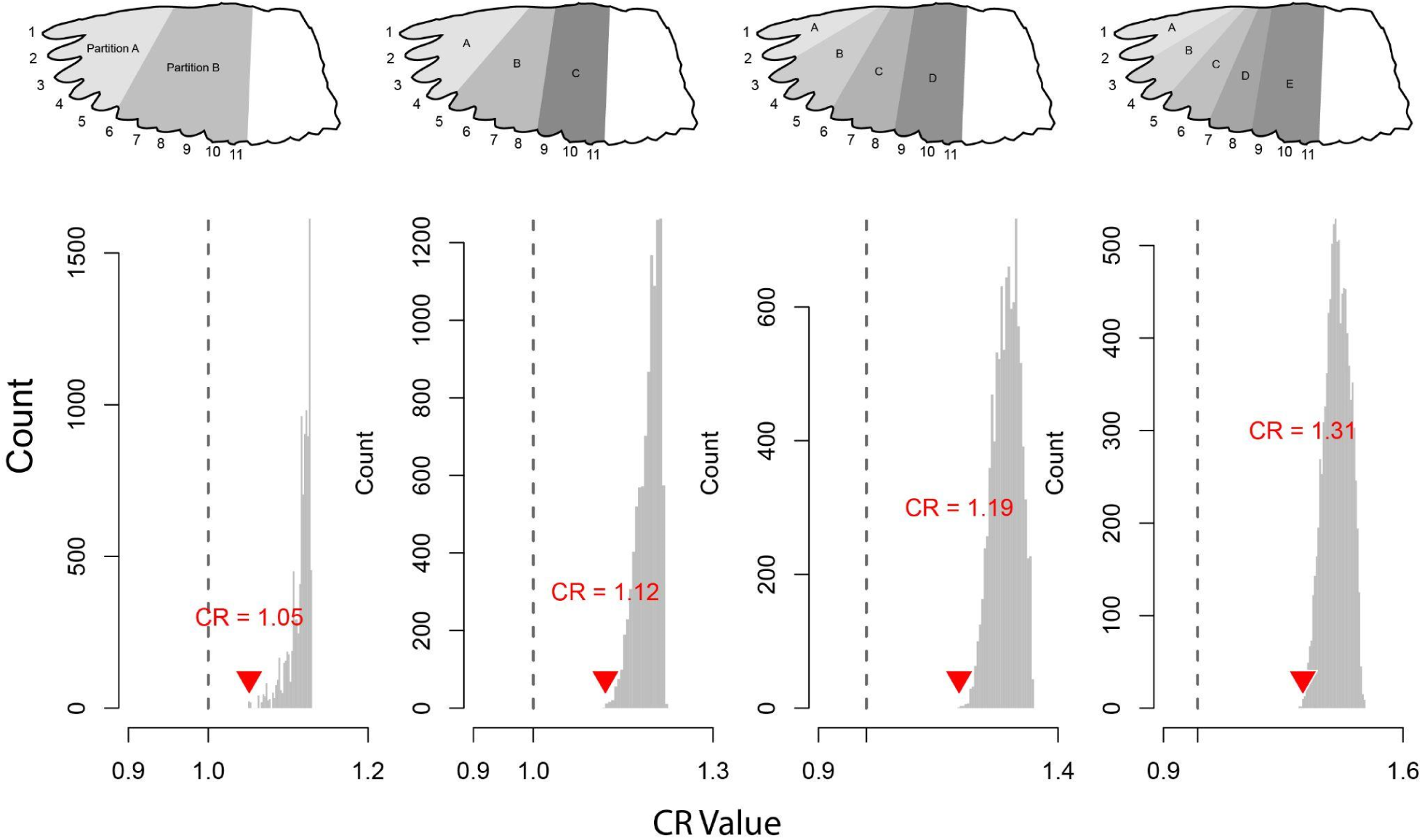
Modularity test of the dimensional primary feathers. Wing illustrations depict schemes of partitioning feathers into hypothetical modules. Modularity was tested using the Covariance Ratio test from Adams (2016). CR < 1.0 indicates modularity, while CR > 1.0 indicates integrated evolution. The recovered CR values for each partitioning scheme are shown with an inverted red triangle. The gray background histograms show the result of a permutation test, randomly shuffling the feathers among the putative modules (*n* = 10,000).

**Figure S3.**
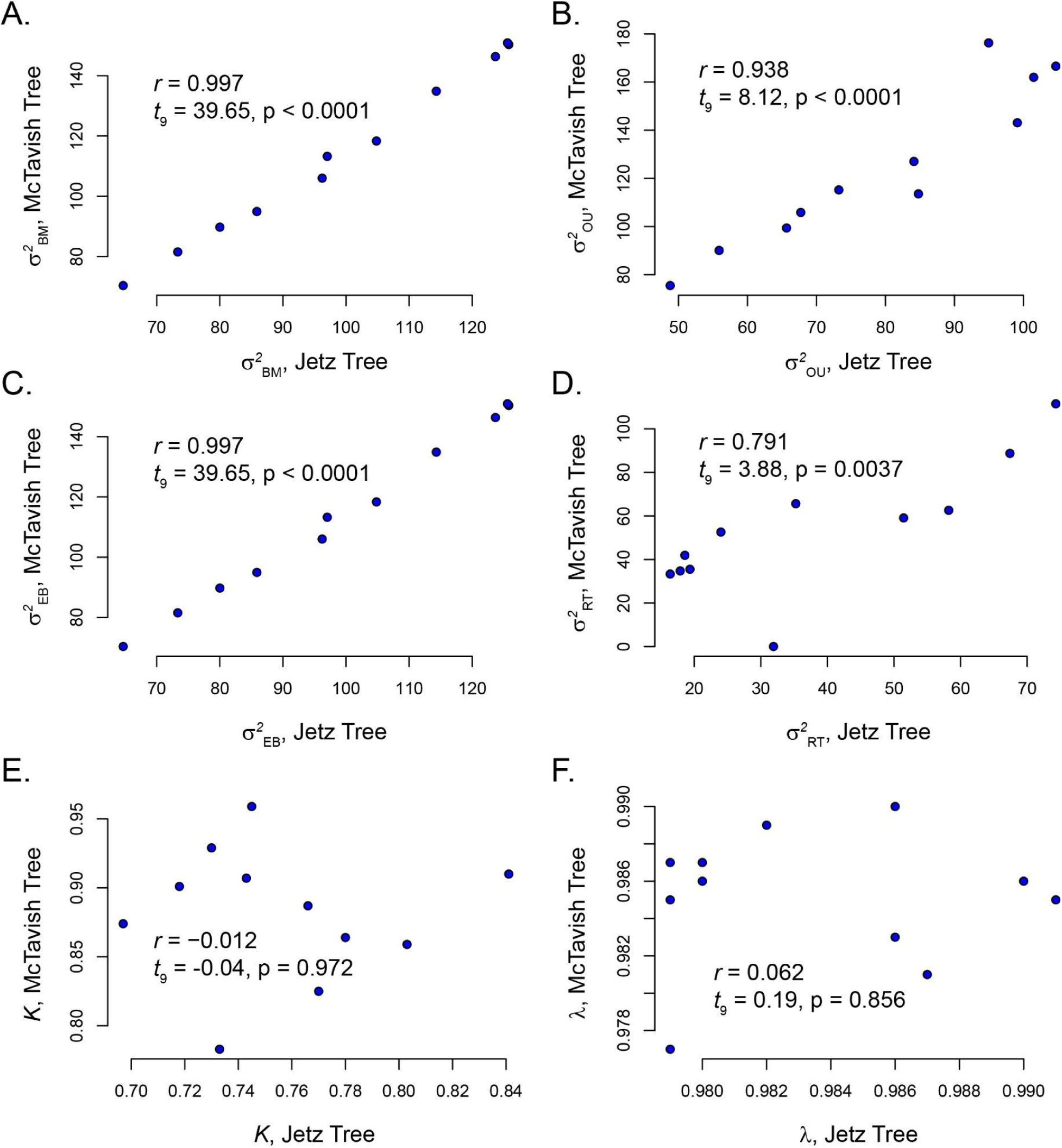
Comparison of results between two phylogenetic reconstructions. Estimates of Evolutionary rate (**σ**^2^) were highly correlated for all four evolutionary models (Brownian motion, BM; Ornstein-Uhlenbeck, OU, Early Burst, EB; Rate Trend, RT) between the Jetz et al. and McTavish et al. phylogenies (A-D). Blomberg’s K (E) and Pagel’s *λ* (F) did not show similar correlations, probably owing to the small variance in estimates on both phylogenies, but the estimated values were qualitatively similar.

